# LOCATOR: feature extraction and spatial analysis of the cancer tissue microenvironment using mass cytometry imaging technologies

**DOI:** 10.1101/2023.03.22.533745

**Authors:** Rezvan Ehsani, Inge Jonassen, Lars A. Akslen, Dimitrios Kleftogiannis

## Abstract

Recent advances in highly multiplexed imaging have provided unprecedented insights into the complex cellular organization of tissues, with many applications in translational medicine. However, downstream analyses of multiplexed imaging data face several technical limitations, and although some computational methods and bioinformatics tools are available, deciphering the complex spatial organisation of cellular ecosystems remains a challenging problem. To mitigate this problem, we develop a novel computational tool, LOCATOR (ana**L**ysis **O**f **CA**ncer **Ti**ssue micr**O**envi**R**onment), for spatial analysis of cancer tissue microenvironments using data acquired from mass cytometry imaging (MCI) technologies. LOCATOR introduces a graph-based representation of tissue images to describe features of the cellular organisation and deploys downstream analysis and visualisation utilities that can be used for data-driven patient risk stratification. Our case studies using MCI data from two well-annotated breast cancer cohorts re-confirmed that the spatial organisation of the tumour-immune microenvironment is strongly associated with the clinical outcome in breast cancer. In addition, we report interesting potential associations between the spatial organization of macrophages and patients’ survival. Our work introduces an automated and versatile analysis tool for MCI data with many applications in future cancer research projects.

Datasets and codes of LOCATOR are publicly available at https://github.com/RezvanEhsani/LOCATOR.

## INTRODUCTION

Mass cytometry imaging (MCI) is becoming an important technology for basic science and clinical research (Giesen et al. 2014). With MCI clinical samples can be simultaneously analysed for up to 40 proteins/markers with spatial resolution, providing unprecedented opportunities for the in-depth study of the histology and pathophysiology of tissues (Milosevic, 2023; Baharlou et al. 2019). So far, the prevalent technologies namely Imaging Mass Cytometry (IMC) and Multiplexed Ion Beam Imaging (MIBI), have been used for biomarker discoveries, exploration of intercellular interactions and investigation of cellular micro-niches focusing on cancer, diabetes, and other complex diseases. Especially in cancer research, the use of MCI provides novel understanding of the heterogeneity of cell phenotypes (Angelo, et al., 2014; Groom, 2019; Helmink, et al., 2020; Yuan et al. 2016, Nascimento, et al., 2022; Fu et al., 2021) within the **<u>tumour-immune microenvironment (TIME)</u>**. TIMEs association with the clinical outcome via MCI has also opened new avenues for personalised diagnosis and treatment (Jackson et al. 2020; Bhate et al. 2022; Chen, et al., 2020; Goltsev, et al., 2018; Cabrita et al., 2020).

Although the MCI data acquisition is conceptually simple, the visualisation, pre-processing and downstream analysis of imaging data face many challenges. Since MCI data are spatial and multiparametric, novel analytical approaches are needed. Milosevic (Milosevic, 2023) reviews many of these and places them into different categories summarised as follows:1) MCI data visualisation and pre-processing including tools and pipelines available such as CellProfiler (Carpenter, et al., 2006), CATALYST (Chevrier, et al., 2018), RedSEA (Bai, et al., 2021), Steinbock, ImcRtools (Windhager, et al., 2021) and MCMICRO (Schapiro, et al., 2022); 2) cell segmentation including the tools DeepCell (Van Valen, et al., 2016), Ilastik (Berg, et al., 2019), MATISSE (Baars, et al., 2021), CellPose (Stringer, et al., 2021); 3) cell phenotyping with tools such as Astir (Geuenich, et al., 2021), CELESTA (Zhang, et al., 2022) and CellSighter (Amitay, et al., 2022) including state-of-the-art unsupervised clustering algorithms such as Phenograph (Levine, et al., 2015) and FlowSOM (Gassen, et al., 2015); and 4) downstream spatial analysis workflows. It should be noted that some tools falling within category 1, also include cell segmentation and phenotyping utilities so that they can also be used for data exploration tasks.

The available tools/methods belonging to category 4 are closer to precision medicine applications, and they can be in principle used to answer important biological and clinical questions utilising the lens of MCI. The tools within category 4 can be divided into two main subgroups: 4a) general spatial analysis toolboxes; and 4b) cancer-specific spatial analysis frameworks. A striking example within subgroup 4a is CytoMAP (Stoltzfus, et al., 2020) that has been used to explore the myeloid cell localization in lymph nodes of murine samples. Another general toolbox for data visualisation with some downstream analysis options available is called ImaCytE (Somarakis et al. 2021). The Giotto (Dries, et al. 2021) toolbox is more appropriate for the analysis and visualization of spatial transcriptomic data. SPF and DenVar (Seal, et al., 2022; Vu, et al., 2022) are “threshold-free” algorithms to stratify tissue images given the expression profile of a functional marker for the purpose of risk assessment, whereas the lisaClust (Patrick, et al., 2023) algorithm can be used to identify distinct cellular regions within tissue images.

Focusing on subgroup 4b, several spatial analysis frameworks have been proposed to associate TIMEs with the clinical outcome of cancer patients. In a seminal study, Keren and colleagues developed a data-driven approach to investigate the spatial proximity of cell types within breast cancer tissues. The algorithm was used to stratify a cohort of 41 triple-negative breast cancer (TNBC) patients and link specific tissue organization patterns with immune regulation and patient survival (Keren, et al., 2018). Other seminal studies in breast cancer utilised spatial community analyses to unravel phenotypic-genotypic associations linked with patients’ clinical outcome (Ali, et al., 2020; Jackson, et al., 2020). A cellular neighbourhood analysis was implemented by Schürch and colleagues to investigate the organization of tumour and immune components in low-versus high-risk colorectal cancer patients (Schürch et al. 2020). Another algorithm for the detection of connected sets of similar cells (referred to as patches) was proposed by Hock and colleagues and used to assess the expression of multiple chemokines in infiltrated tumours of melanoma patients (Hoch et al. 2022). Finally, Danenberg et al. performed a large-scale analysis of MCI data (>690 samples) from breast tumours and demonstrated the importance of “connectivity features” within TIMEs to link pheno-genomic characteristics with clinically relevant subgroups (Danenberg, et al., 2022).

Most of the previous computational studies, although very influential within the cancer research field, have some technical shortcomings or limitations. Many of the developed frameworks are neither automated nor open source hindering effective re-use and reproducibility. Also, there is a lack of general cancer research tools enabling association of TIME patterns with clinicopathological characteristics of patients in a flexible/modular manner. The existing algorithms are mainly designed to answer study-specific questions, and thus they cannot be readily extended or generalized to investigate the role of arbitrary cell types of interest within TIMEs. Moreover, the lack of a standardized mathematical model to describe cell-to-cell relationships, makes it challenging to generalize findings and compare results from different studies, potentially also utilizing different MCI technologies. Finally, the existing methodologies cannot be readily used to engineer numerical variables (i.e., features) of TIMEs, that can subsequently be fed to machine learning algorithms for improved patient risk stratification. Supplementary Table 1 provides an overview of the most relevant methods. Altogether, we conclude that there is an increasing need for the development of cancer research tools to facilitate hypothesis testing and data-driven MCI research in the new era of precision cancer medicine.

To mitigate these limitations, we introduce LOCATOR (ana**L**ysis **O**f **CA**ncer **Ti**ssue micr**O**envi**R**onment), a novel spatial analysis tool for MCI data from cancer patients. LOCATOR is implemented in the R language that is widely used in biomedical research, and it is specifically designed to engineer features describing local/global cellular composition of tissues, as well as structural features of cells and their interactions within TIMEs. To achieve this goal, LOCATOR proposes a graph-based feature engineering approach. Importantly, LOCATOR is versatile, independent of the imaging technology used (e.g., IMC or MIBI), and can be applied to identify possible associations between TIMEs and other molecular and clinical variables.

We use LOCATOR to analyse publicly available data from two breast cancer cohorts as well as synthetic data. Our results corroborate previous findings about the role of TIMEs, highlighting LOCATOR’s ability for TIME-driven patient risk stratification. With LOCATOR we also investigate novel multicellular features of macrophages and report potentially interesting associations with distinct clinicopathological parameters. We anticipate that LOCATOR could find many useful applications in future cancer research projects.

## RESULTS

### LOCATOR utilizes a graph-based representation of cellular ecosystems for feature discoveries

The tumour microenvironment is composed of different cell types that form spatially organized cellular ecosystems. Within these cellular neighbourhoods immune and tumour cells interact directly or indirectly with each other adopting different mechanisms and functions, and many studies have emphasized the role of TIMEs in suppressing or promoting tumour growth (Biswas, et al., 2022; Fu, et al., 2021; Hsieh, et al., 2022; Yuan, 2016). With this idea in mind and to be able to easily data-drive research towards better patient stratification, we present LOCATOR. LOCATOR utilises the spatial location (i.e., X and Y coordinates in 2D) of cells to generate local cellular neighbourhoods in the tissue microenvironments using a user-defined radius. LOCATOR’s implementation is “flexible” since different investigators may use it appropriately to dissect cellular neighbourhoods screened with any of the prevalent MCI technologies. To gain maximised information about each individual cellular neighbourhood and subsequently generate comprehensive spatial features (i.e., numerical variables) describing the interactions between cells, LOCATOR adapts a graph-based representation. With this abstract formulation, all cells in a cellular neighbourhood are modelled as vertices and links between them are represented as edges (i.e., cell-cell interactions). LOCATOR defines and computes three different categories of features: 1) cellular abundance (%) in the neighbourhood; 2) features describing the connectivity of cell types in the neighbourhood; 3) features described by standard graph theory metrics and graph algorithms. A more detailed technical overview of the feature matrix generated by LOCATOR is described in the Materials and Methods section. Figure 1 shows a blueprint of LOCATOR’s feature engineering component.

**Figure 1:**
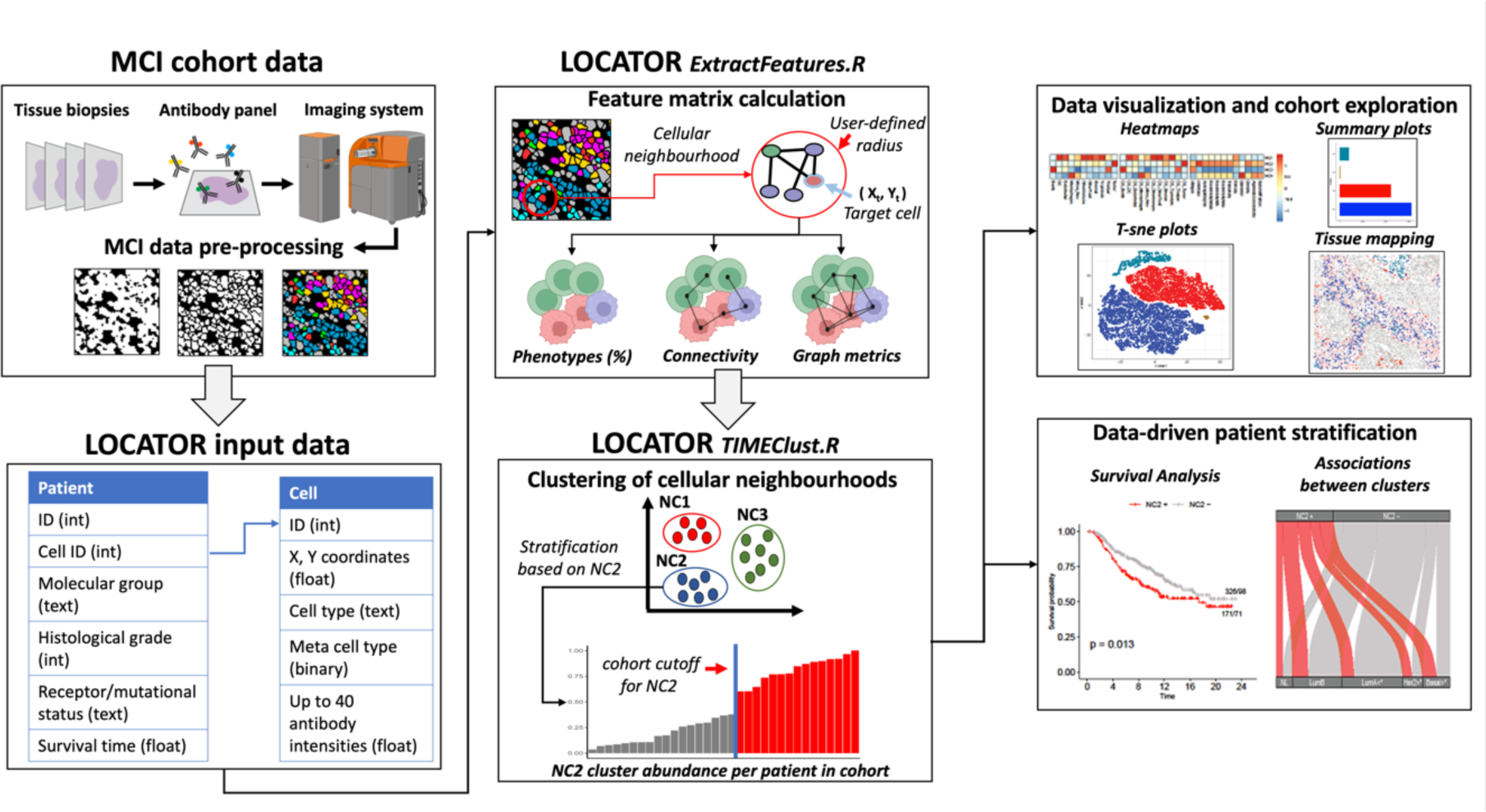
LOCATOR workflow. The flowchart describes the data acquisition and pre-processing process of multiplexed imaging data that is required to generate the LOCATOR’s input data. LOCATOR workflow uses two programs written in R language namely, ExtractFeatures and TIMEClust. ExtractFeatures is used to define cellular neighbourhoods and generate graphs. These graphs are used for feature extraction. Next, TIMEClust program uses these features to cluster the cellular neighbourhoods of interest. At the patient level, patients as enriched or decreased for each of the identified clusters. This provides opportunities for MCI data visualisation and downstream analyses for data-driven patient stratification tasks. (image created with BioRender.com)

After the feature generation step, LOCATOR deploys an unsupervised consensus clustering approach to group together cellular neighbourhoods with similar/dissimilar spatial properties as described by numerical TIME features. For each of the identified cellular clusters, patients can be enriched or depleted by comparing patient-specific values to the mean/median cluster values computed at the cohort level. This information can be directly used to infer associations between TIMEs and patient-specific clinicopathological parameters. The proposed data engineering approach is quite versatile enabling effective downstream analyses, in combination with artificial intelligence algorithms or any other data science technique. A step-by-step description of LOCATOR’s workflow is available in the Supplementary Figure 1.

### LOCATOR systematises TIME-driven patient stratification in breast cancer

To showcase the use of LOCATOR for MCI data exploration and TIME-driven patient stratification, we performed two case studies using as input data from two independent breast cancer studies (Danenberg, et al., 2022; Keren, et al., 2018). Since one of the most promising clinical applications of MCI is to investigate the role of immune cells in the cellular microenvironment, our case studies are anchored around immune cell types, with a primary focus on macrophages.

Macrophages are innate immune cells with phagocytic function and heterogeneous characteristics (Qiu, et al., 2018). In several breast cancer studies, high infiltration of tumour-associated macrophages (TAM) is associated with shorter patient survival, but unfortunately, our knowledge about their role and regulation is rather limited (Choi, et al., 2018; Kumari and Choi, 2022; Lin, et al., 2019). For extra experimentation with LOCATOR, we also provide computational analysis results for another immune cell type, the T cytotoxic cells, with known (Dushyanthen, et al., 2015; Stanton and Disis, 2016; Li, et al., 2021) prognostic and predictive value in specific breast cancer subgroups (HER2+, and Triple-negative).

To explore LOCATOR’s capabilities, we compared its results to independent clinicopathological information of patients such as survival time, PAM50 molecular subtype, histological grade, and hormone receptor status when available (Supplementary Figure 2a, 2b). Next, LOCATOR’s results are compared with computationally derived results published independently by Keren et al. and Danenberg et al.

#### 1. Case study using Keren dataset

We apply LOCATOR to analyse 33 TNBC samples available in the Keren dataset. From the original 41 samples of this dataset, five samples are “cold” tumours (i.e., very low immune presence) and they are filtered out. We also filter out three samples with less than 20 macrophages to ensure robust feature matrices calculations for all samples included in the study. Note that this parameter can be customized by the users depending on the study design and the available cells of interest. LOCATOR’s consensus clustering approach identifies four cellular clusters of macrophages (NC1, NC2, NC3 and NC4), and the mean abundance at the cohort level is used as a cut-off to identify patients enriched and depleted for each of these clusters. Figure 2 shows example images produced using LOCATOR’s API, including a heatmap representation of TIME features, summary plots focusing on the cellular neighbourhoods surrounding macrophages, a t-distributed stochastic neighbour embedding (t-SNE) of the clustering results and an example of tissue reconstruction mappings with clustering annotations. Similar outputs focusing on cellular neighbourhoods surrounding T cytotoxic cells are shown in Supplementary Figure 3.

**Figure 2.**
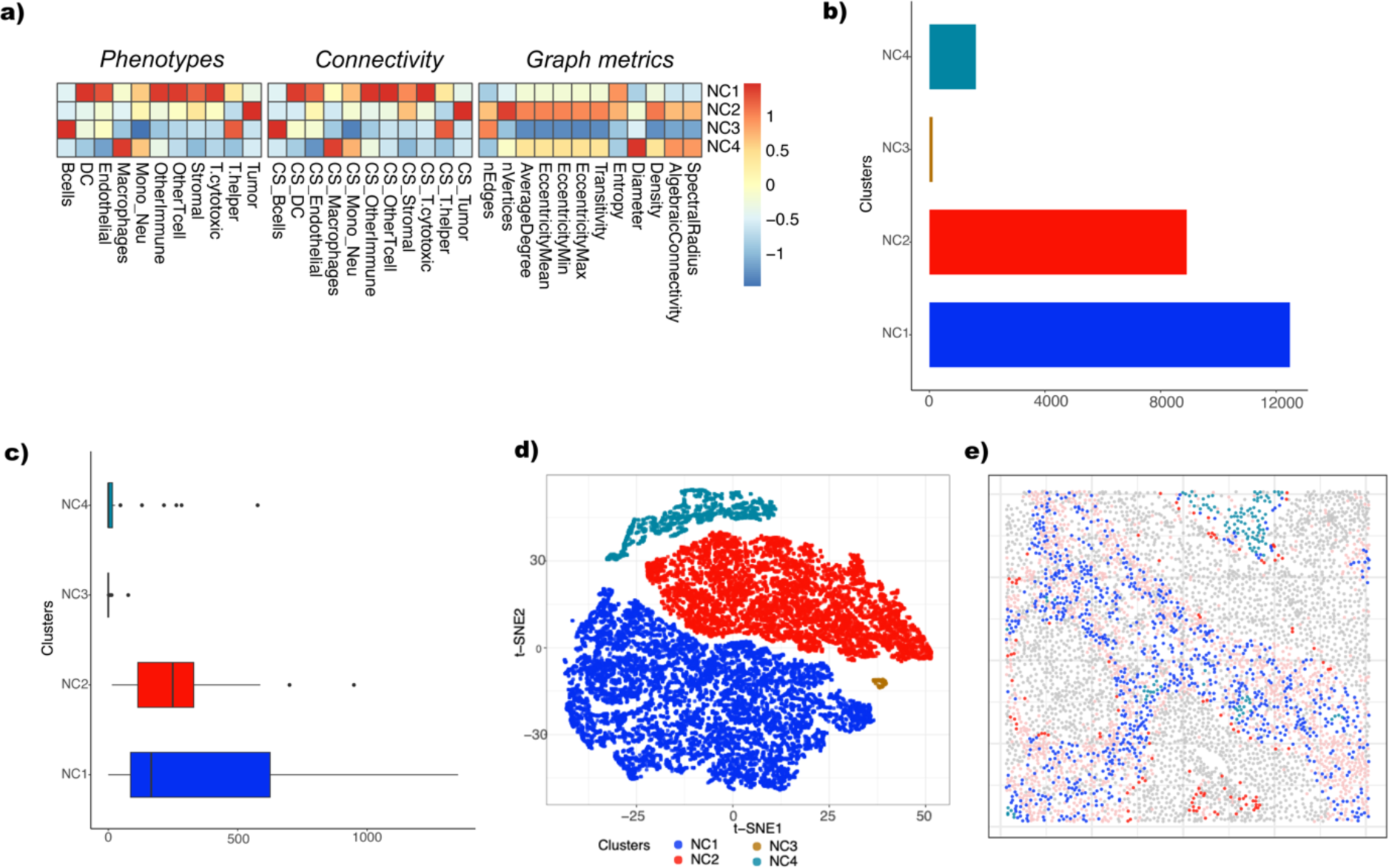
LOCATOR outputs for Keren dataset focusing on macrophage cell type. **a) Heatmap representation of the feature vector for the identified clusters.** b) Bar plot showing cellular abundance of the identified cellular clusters. c) Boxplot showing the number of macrophages per patients across different cellular clusters. d)t-SNE visualisation of the identified cellular clusters. e) Example of tissue mapping with cellular cluster annotation from one patient (patient No 5). Gray colour indicates tumour cells and pink colour indicates non-tumour cells. All other colours in the image correspond to the detected cellular clusters from subplots c and d.

##### 1.1 Associating LOCATOR’s clustering results with patient survival

Based on LOCATOR’s consensus clustering results for macrophages we identify potential associations with disease outcome. Since survival information is independent from the data generation and clustering processes used by LOCATOR, it suggests that the spatial patterns found by LOCATOR have relevance for disease development and outcome. Using cellular clustering results at the patient level (i.e., number of patients enriched or depleted for each of NC1-NC4 clusters) we perform univariate Kaplan-Meier survival analysis. Figure 3a shows the survival analysis results for cluster NC2. We observe that patients enriched for this cluster (called NC2+) achieve worse survival compared to the depleted (NC2-) patients (long-rank p-value=0.018). Upon inspection of the connectivity features (Figure 2a), we observe that cluster NC2 has higher feature values for tumour-macrophages connectivity scores, whereas from the graph metric-based features the number of vertices is also high. Such findings might be linked to the underlying biology of tumour-associated macrophages within TIMEs, opening new avenues for deciphering complex mechanisms of macrophage activation/regulation.

**Figure 3.**
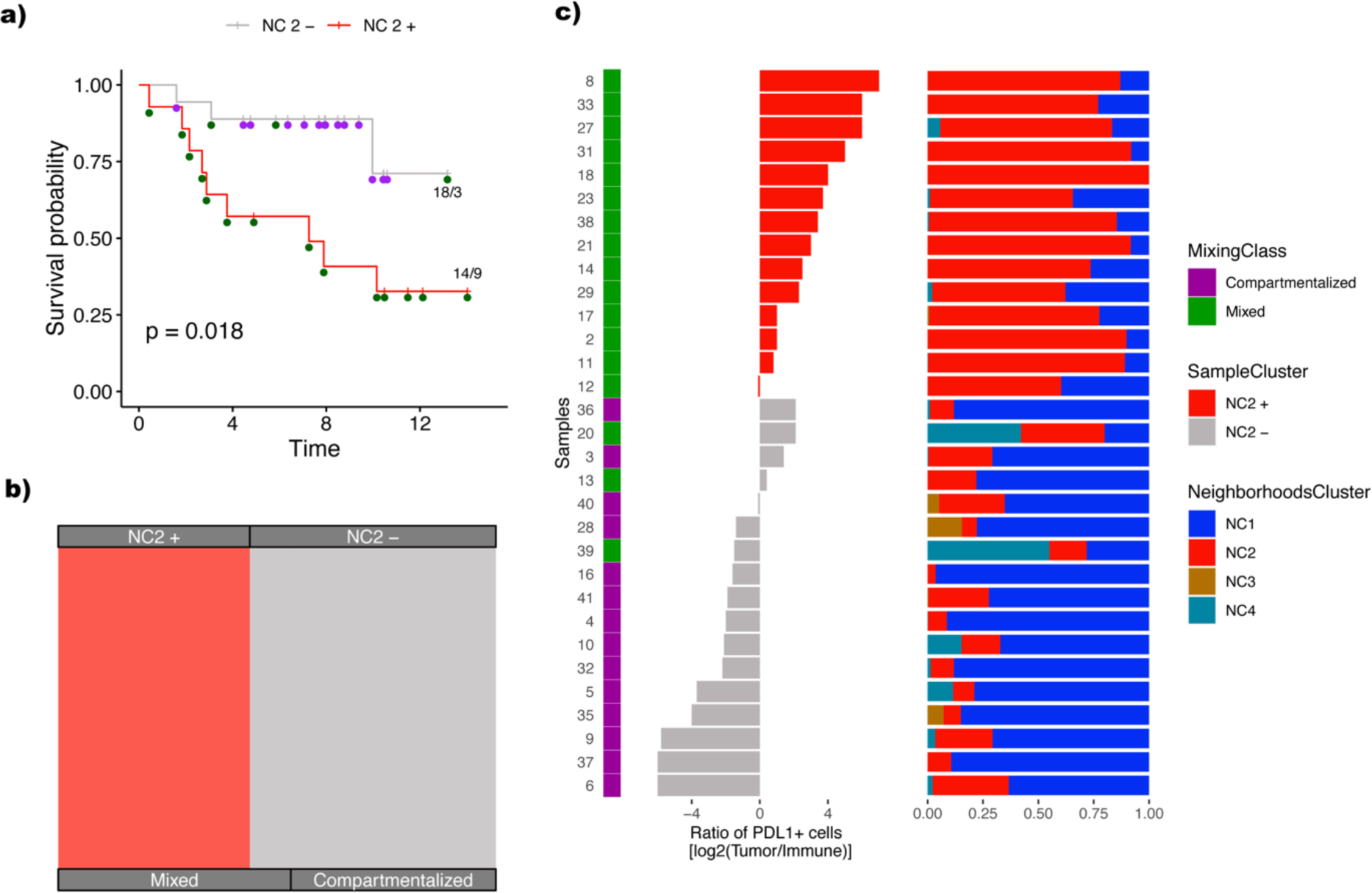
Case study 1: Association of the identified NC2 cellular cluster with clinicopathological parameters and independent published results. a) Survival plot where patients are classified as enriched (NC2+) or depleted (NC2-) based on their mean abundance for NC2. b) Alluvial plot showing the fraction of NC2 enriched/depleted patients and their underlying tissue mixing classification. c) From left to right, original classification of patients and PDL1 score from Keren et al., where colour code is based on the patients enriched/decreased for NC2 cellular cluster. The stack bar shows the abundance of different macrophage cellular clusters identified by TIMEClust across all patients.

##### 1.2 Comparison with the published mixing score for tissue classification

Next, we compare our clustering results using the mixed and compartmentalized tissue annotation derived from the mixing score deployed by Keren and colleagues. Figure 3b shows that patients of the NC2+ subgroup belong mainly to the mixed tissue type, which is linked with poorer survival, whereas patients of NC2-subgroup have mainly compartmentalized tissue types and better survival.

Further we compare LOCATOR’s NC2 patient subgroups with the PDL1 immunoregulatory protein score published by Keren and colleagues. In the original study by Keren et al. higher PDL1 score levels are associated with poorer survival, mixed tissue type and cancer progression. Our results (Figure 3c) focusing on NC2 cellular cluster of macrophages corroborate all the previous findings about PDL1 levels and the underlying tissue architecture.

#### 2. Case study using Danenberg dataset

We also apply LOCATOR to analyse 497 cases from the Danenberg dataset. From the original 693 cases of this dataset, we filter out 196 samples with less than 20 macrophages to ensure robust feature matrix calculation. Focusing on macrophages, LOCATOR’s consensus clustering approach identifies 14 cellular clusters (NC1 to NC14). We use the mean abundance at the cohort level as a cut-off to determine enriched and depleted patients for each of these cellular clusters. Figure 4 shows example images that can be produced using LOCATOR’s visualisation functions for this dataset. Similar outputs focusing on cellular neighbourhoods surrounding T cytotoxic cells for this dataset are shown in Supplementary Figure 4.

**Figure 4.**
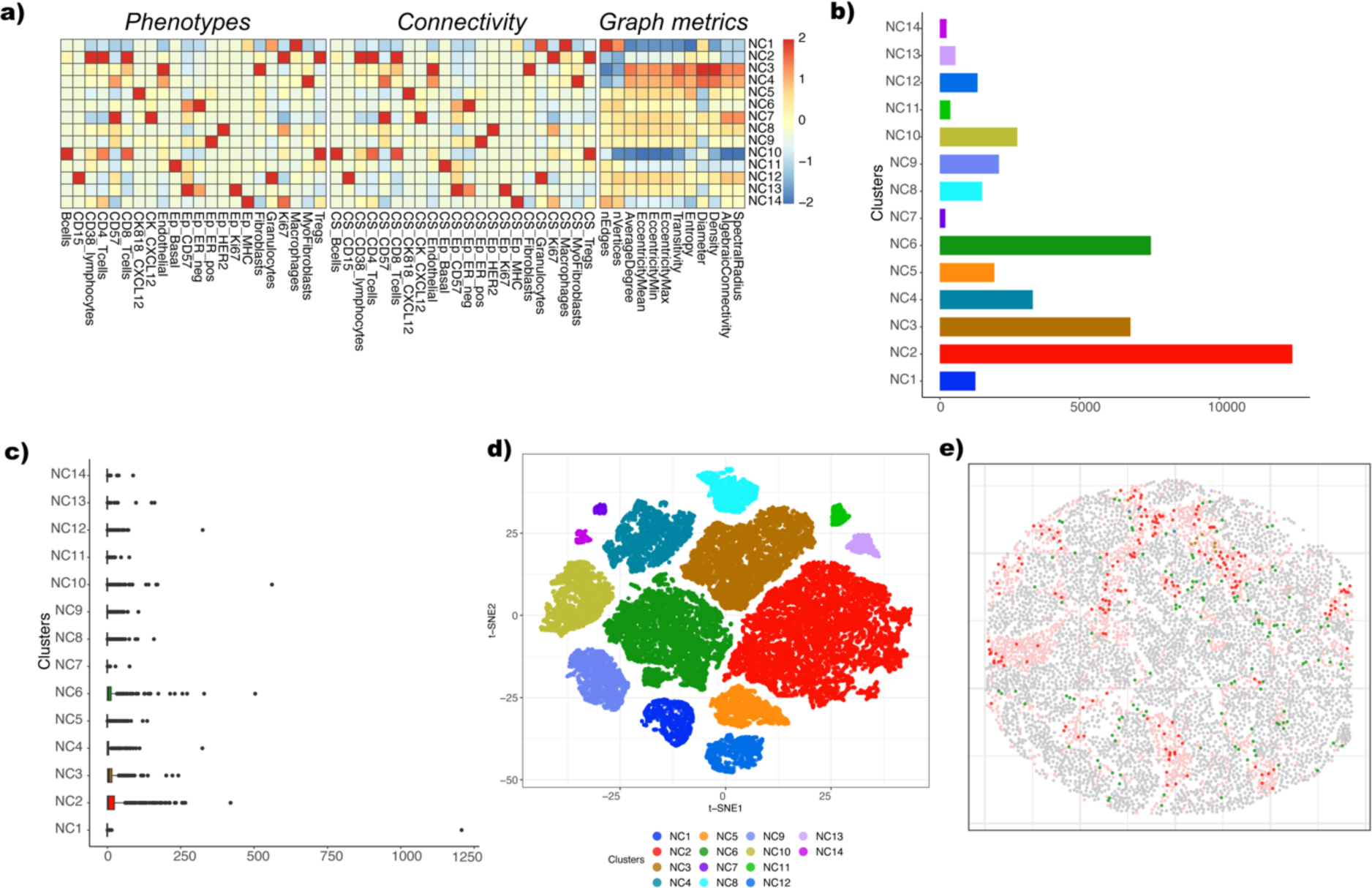
LOCATOR outputs for Danenberg dataset focusing on macrophage cell type. a) Heatmap representation of the feature vector for the identified clusters. b) Bar plot showing cellular abundance of the identified cellular clusters. c) Boxplot showing the number of macrophages per patients across different cellular clusters. d)t-SNE visualisation of the identified cellular clusters. e) Example of tissue mapping with cellular cluster annotation from one patient (patient No 191). Gray colour indicates tumour cells and pink colour indicates non-tumour cells. All other colours in the image correspond to the detected cellular clusters from subplots c and d.

##### 2.1 Associating LOCATOR’s clustering results with patient survival and other clinical information

Using LOCATOR, we investigate whether the acquired cellular clustering results are associated with patient survival and other independent clinical information namely PAM50, histological grade, and hormone receptor status. Based on univariate Kaplan-Meier survival analysis we find that patients enriched for NC2 (NC2+) and NC8 (NC8+) cellular clusters are linked with shorter survival whereas patients enriched for cellular cluster NC3 (NC3+) are associated with longer survival (Figure 5 a, b, and c).

**Figure 5.**
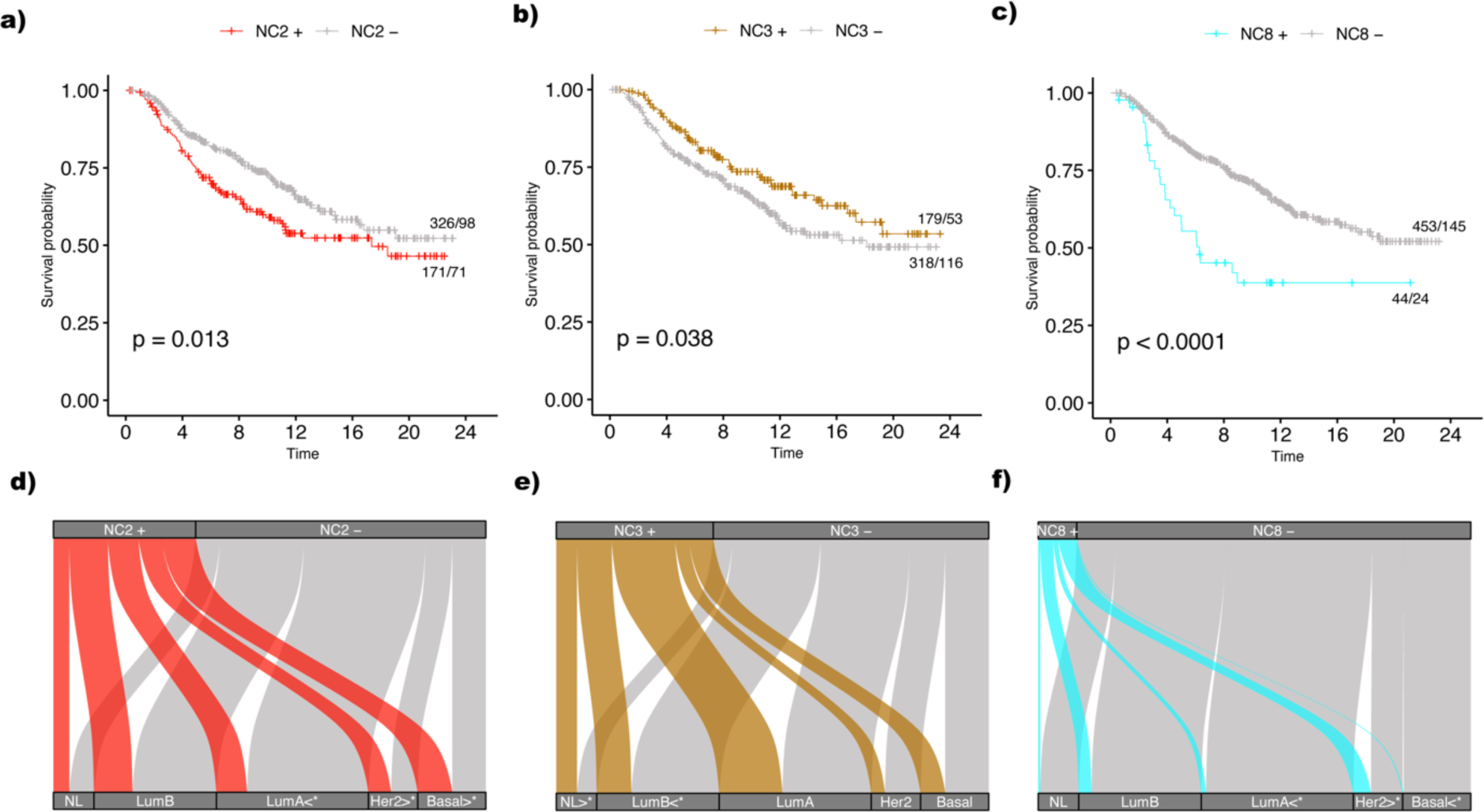
Case study 2: Association of NC2, NC3 and NC8 identified cellular clusters with patient survival and PAM50 classification. a-c) Survival plots where patients are classified as enriched (NC2+, NC3+, NC8+) or depleted (NC2-, NC3-, NC8-) based on their mean cellular abundances. d-f) Alluvial plots showing the fraction of NC2, NC3 and NC8 enriched/depleted patients and their underlying PAM50 classification. Significance of the results is obtained using Fishers-exact test at a p-value levels of 0.05 (results marked with *, and Supplementary Figure 7 for details).

Based on PAM50 criteria, we find that NC2+ patient subgroup includes mainly Her2+ and Basal-like cases, whereas the number of LumA cases, which are in principle associated with favourable outcome, is proportionally smaller compared to NC2-patient subgroup. Based on histological grade information, we observe that the NC2+ subgroup includes more Grade 3 cases compared to NC2-. Using ER hormone receptor status, we also see more ER-cases for the NC2+ patient subgroup compared to NC2-.

Based on histological grade information, we find that NC3+ patient subgroup includes proportionally smaller number of Grade 3 cases compared to NC3-, and as expected NC3+ contains mainly patients with Grade 1. Based on ER hormone receptor status, NC3+ patients include less cases with ER- and more ER+ cases compared to NC3-.

Focusing on PAM50 criteria, for cluster NC8+ we observe proportionally smaller number of LumA cases and more cases with Her2 compared to NC8-. According to histological grade, NC8+ patient subgroup includes more Grade 3 cases (Supplementary Figure 5) compared to NC8-. At last, based on ER hormone receptor status, we observe that NC8+ patients are in their majority ER-cases. Supplementary Figure 4 shows additional clustering analysis results focusing on T cytotoxic cells. We emphasise on three clusters (Supplementary Figure 6), namely NC2, NC3 and NC4, that are potentially linked with clinically relevant features of TIMEs. Specifically, patient subgroups NC2+, NC4+ and NC3-that achieve shorter survival are enriched for some clinical features of “aggressive cancers” namely basal-like PAM50 phenotypes, ER negativity and higher tumour grade.

##### 2.2 Comparison with published clustering results

We also compare LOCATOR’s clustering results with the clustering results reported by Danenberg and colleagues (Figure 5, Supplementary Table 2, and Supplementary Figure 5). Focusing on NC2+ patient subgroup that achieve shorter survival, we find that this cluster includes an increased number of cases from IntClust 4- and IntClust 10 reported by Danenberg and colleagues compared to NC2-patient subgroup. Both IntClust 4- and IntClust 10 are linked to poor prognosis in the original publication. In NC2+ patient subgroup we also find proportionally less cases from IntClust 8 and IntClust 7 clusters reported by Denenberg et al. compared to NC2-patient subgroup. Note that both IntClust 8 and IntClust 7 clusters are linked with good prognosis based on the original published results.

Focusing on NC3+ patient subgroup that achieve longer survival compared to NC3-, we find (Figure 5, Supplementary Table 2, and Supplementary Figure 5) that NC3+ includes proportionally more Normal Like cases.

Finally, for NC8+ patient subgroup that achieve shorter survival compared to NC8-, we observe that NC8+ includes mostly cases from IntClust 5-cluster of the published results that is linked with poor survival, whereas it includes proportionally smaller number of cases from IntClust 3, IntClust 7, and IntClust 8 clusters, all associated with good survival based on the original results published by Danenberg and colleagues.

Taken together, the results of both case studies with a primary focus on macrophages, corroborate findings from independent publications/analyses (Keren et al. 2018 and Danenberg et al. 2022). Our results highlight LOCATOR’s ability to test hypotheses and discover novel TIME-dependent patient subgroups that can be used for effective patient stratification in subsequent cancer research projects.

### Investigating the potential impact of neighbourhood size for TIME-driven patient stratification

The ability to investigate accurately cellular microenvironments via MCI depends strongly on which value is chosen for the neighbourhood size parameter (i.e., radius). For example, “over-binning” cells into very large cellular neighbourhoods might lead to loss of important biological signals. Conversely, reducing the neighbourhood size might generate many and noisy small networks that might not be biologically and clinically relevant. The selection of appropriate neighbourhood size is also important for practical reasons: it affects the complexity of the computational analysis and subsequently the running time of the tool and the required computational resources. Existing tools such as CytoMAP and the analysis frameworks implemented by Keren et al., Jackson et al. and Danenberg et al. selected the neighbourhood size after empirical assessment leading to a radius of 100 pixels for the Keren dataset (MIBI) and of 50 pixels for the Danenberg dataset respectively (IMC Hyperion). The same neighbourhood sizes were used by LOCATOR in the case studies described above.

Here, to systematically assess the potential impact of neighbourhood size for TIME-driven patient stratification by LOCATOR’s TIMEClust program, we perform additional analyses. Using the Keren dataset as a benchmark, we repeat the clustering step with neighbourhood sizes [50, 75, 100,125,150]. For every run we focus on the cluster that achieves the lowest log-rank p-value of Kaplan-Meier survival analysis, assuming that this is the cluster most closely linked to the clinical outcome. We use the mixed and compartmentalised tissue types reported by Keren and colleagues, and we estimate Accuracy, Sensitivity, Specificity using known formulas (Soufan, et al., 2015). Figure 6a conveniency demonstrates that the neighbourhood size does indeed matter, with the value of 50 achieving the lowest Specificity. Interestingly, sizes between 75 and 125 achieve very comparable results with size of 75 being the best in terms of Accuracy and Specificity, but the Kaplan-Meier survival results did not reach statistically significant levels (Figure 6b, p value threshold of 0.05). In summary, our experimentation indicates that size of 100 is indeed a very reasonable selection for the Keren dataset (MIBI).

**Figure 6:**
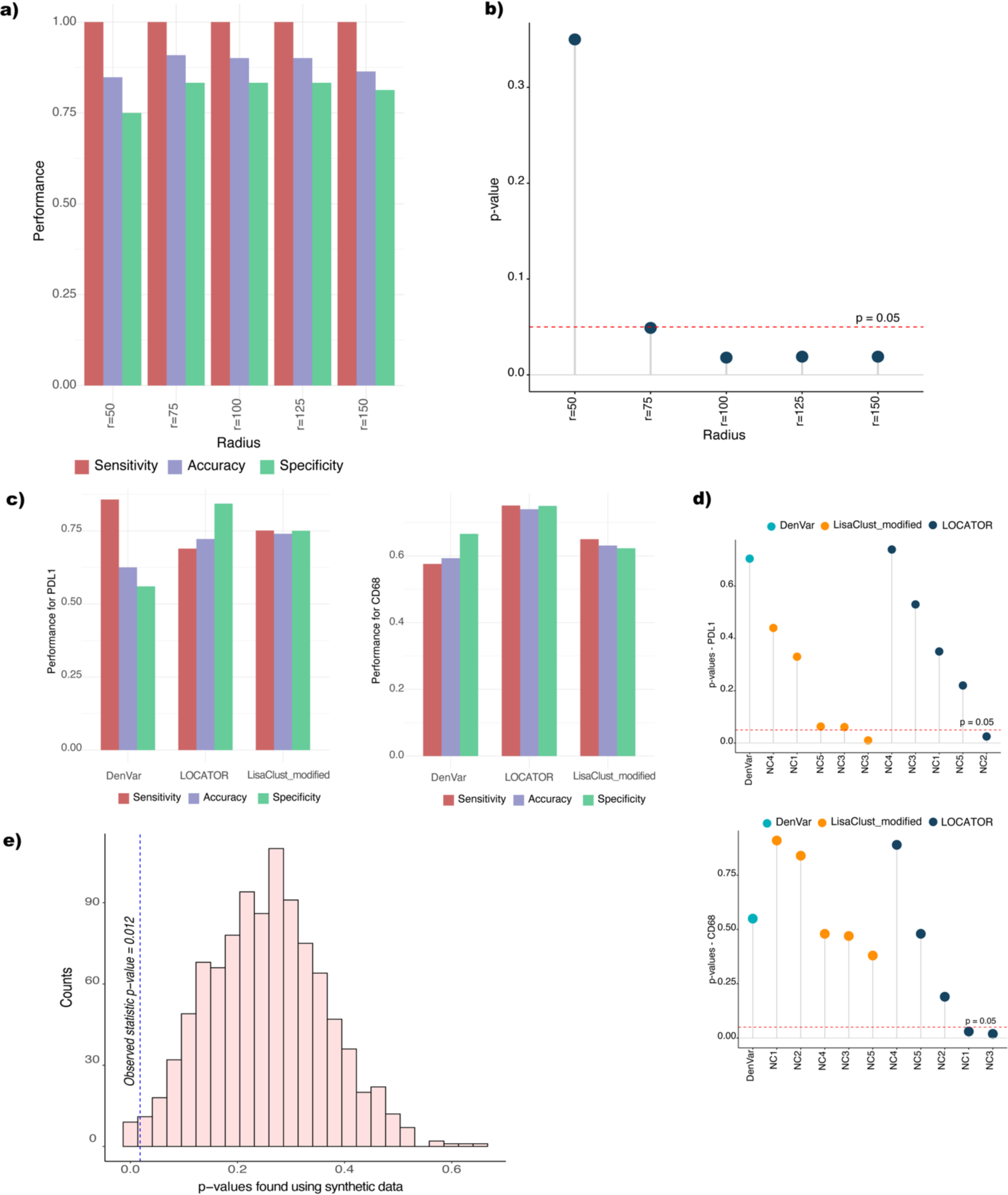
LOCATOR’s performance evaluation. a) Barplot showing Sensitivity, Specificity and Accuracy of LOCATOR’s results for different values of Radius parameter. Tissue annotations from Keren dataset are used as a benchmark. b) Dot plot summarising the survival analysis (log-rank p-values) using the clustering results from (a). All p-values were obtained using univariate Kaplan-Meier method, considering a p-value cutoff of 0.05. c) Comparison analysis results between DenVar, LisaClust and LOCATOR. We report average Sensitivity, Specificity and Accuracy, using as benchmark tissue annotations from Keren dataset. d) Dot plot summarising the survival analysis (log-rank p-values) using the clustering results from (c). All p-values were obtained using univariate Kaplan-Meier method, considering a p-value cutoff of 0.05. e) Assessing the significance of LOCATOR’s survival analysis results using as input synthetic data.

### Comparing LOCATOR with independent spatial analysis methods

We compare LOCATOR’s clustering results at the patient level with two alternative spatial analysis methods namely DenVar and lisaClust. As a benchmark we focus on Keren dataset, and we consider the mixed and compartmentalised tissue annotations. Since DenVar requires as input the expression of a single marker, we run LOCATOR with input cell annotations based on the scaled expression of PDL1 and CD68 markers (i.e., cells with expression greater/lower than z-score 0.5). To facilitate fair assessment of the competitor methods, we also run lisaClust using as input the same set of positive/negative cells for PDL1 and CD68 markers. However, lisaClust’s original implementation cannot be used for patient stratification. To mitigate this limitation, we deploy our own version of lisaClust that enables patient stratification called lisaClust_modified. In this version, the outputs returned by lisaClust algorithm are used to identify subgroups of patients depleted/enriched for LISA outputs following the same methodology as in LOCATOR’s TimeClust program.

Figure 6c shows the average Sensitivity, Specificity and Accuracy of the competitor methods (see Supplementary Table 3 for the complete set of results). We observe that in most cases LOCATOR achieves the best results on average, whereas using PDL1 lisaClust_modified achieves slightly better Accuracy and Sensitivity. Next, for the best cluster in terms of Accuracy, we perform Kaplan-Meier survival analysis. Interestingly, (Figure 6d) using PDL1 LOCATOR and LisaClust_modified achieved statistically significant results for only one of the detected clusters (NC2 and NC3, p-values = 0.02 and 0.01 respectively). When considering CD68 marker as input, only LOCATOR was able to stratify patients into subgroups of different survival (both NC1 and NC3 clusters with p-values = 0.03, 0.02 respectively).

### Experimentation with synthetic data

In this subsection, we perform a self-consistency test using as input synthetic data. Generating synthetic imaging data is not a straightforward task, as our knowledge about “aggressive” spatial features of cancer is rather limited. Here, to mitigate this limitation, we follow the complete spatial randomness assumption (CRS). Based on CRS cells do not have any preference for any spatial location and connection with other cell types (Stoltzfus et al. 2020). However, since there is increasing evidence that the cellular abundance and the ratio of cancer and immune cells (i.e., infiltration) are clinically relevant features of breast cancer, we maintain the cellular abundances from the Keren dataset as presented in Supplementary Figure 2c. Thus, using as “seed” real cellular abundance distributions we generate a more realistic synthetic MCI cohort with random spatial locations and connections between cells.

Using this synthetic cohort, we repeat LOCATOR’s analysis 1000 times, and for the resulting clusters we perform univariate Kaplan-Meier survival analysis. In this way, it is possible to compute the number of times a statistic (i.e., log rank p value) smaller than that found for the Keren dataset itself, is obseerved. Figure 6e summarises the original statistic and the statistics found using the experimentation with simulated data. The results of Figure 6e are translated to a p value of 0.012, which is significant at a 0.05 threshold. In summary, our self-consistency test conveniently highlights LOCATOR’s power to produce reliable and robust clustering results at the patient level.

## MATERIALS AND METHODS

### LOCATOR overview

#### Required input data

LOCATOR consists of the two programs *ExtractFeatures* and *TIMEClust,* both written in the R language containing functions respectively for feature calculation and for downstream analysis including TIME-driven patient stratification. The method requires the availability of data for a cohort of patient samples screened with any of the available tissue imaging technologies using an antibody panel. LOCATOR can be applied after the raw images have been externally pre-processed, segmented and cells phenotyped. For tissue image segmentation we recommend any of the available state-of-the-art frameworks DeepCell (Bannon, et al., 2021), or ImcSegmentationPipeline (Vito RT Zanotelli, 2022). Once the segmented images have been produced, cell phenotyping can be performed using any of the available state-of-the-art approaches (Milosevic, 2023). A data schema of the required input data is shown in Figure 1. Technically the input data can be provided in an aggregated tabular file (e.g., csv) where each patient is indexed by a unique ID, followed by other relevant cohort-specific clinical information such as molecular subtype (e.g., PAM50 in breast cancer), histological grade, mutational status or hormone receptor status and time of death/relapse. We note that LOCATOR’s software implementation also supports SingleCellExperiment R structures. Each patient’s tissue image contains individual cells indexed by their unique ID, X and Y coordinates. Depending on the antibody panel design, every individual cell is associated with a vector of up to 40 antibody intensities. Every cell is also assigned to a unique phenotype (cell type) returned during the pre-processing step.

Once LOCATOR’s required input data have been loaded, the user must select the target cell type from the available “canonical” phenotypes (e.g., T-cells, B-cells, Macrophages etc) identified during the cell phenotyping step. In addition, with LOCATOR it is also possible for users to select “custom” subsets of cells based for example on the expression of a single functional marker (e.g., all cells enriched/depleted for PDL1 see section “Comparing LOCATOR with independent spatial analysis methods”).

#### Feature discovery

LOCATOR uses the program *ExtractFeatures* to engineer features of TIMEs. LOCATOR deploys a graph-based representation of cellular neighbourhoods to engineer features as follows: for every target cell LOCATOR creates a circular neighbourhood using a predefined radius centered around the cell of interest using the X and Y coordinates. Choosing an appropriate value for the radius depends on the resolution of the imaging technology used. After experimentation (see Results section) we recommend radius=100 pixels for MIBI and radius=50 for Hyperion IMC technologies. Within each cellular neighbourhood LOCATOR constructs an individual undirected graph where cells are vertices and interactions between them are edges. To identify all possible cell-to-cell interactions within a neighbourhood, the Voronoi algorithm is used (Aurenhammer, 1991). To speed this process up, the Voronoi algorithm considers only the cells within the neighbourhood, neglecting possible interactions of cells between different neighbourhoods (e.g., circles that are far apart). Based on this approach, three different categories of features are computed for every neighbourhood/graph (i.e., number of neighbourhoods equals the number of cells of interest) to capture phenotypic, functional, and structural information.

1. ***Cellular abundance (%)*** within the neighbourhood/graph: where the number of features equals the number of phenotypes found in the dataset.
2. ***Connectivity score* (CS)**: a weighted score reflecting the number of interactions and their importance within the neighbourhood. For each cell type within the neighbourhood, we first compute the number of interactions found with all other cell types (i.e., number of edges). Then each edge is weighted with *w1* if both vertexes (cells) are of the same cell type, with *w2* if the vertices are of different cell types but they are both non-tumour cells or both tumour cells, and with *w3* if the vertices are of different cell type but one is non-tumour cell and the other is tumour cell. Default weight parameters are *w1=1, w2=0.5 and w3=0.1*, but this can be adjusted by the user. The final connectivity score for each cell type within a neighbourhood, is computed by summing up all partial weighted scores.
3. ***Graph theory metrics***: for every neighbourhood/graph we compute the number of vertices, the number of edges, average vertex degree, entropy, density, diameter, eccentricity (max, min, mean), transitivity, algebraic connectivity, and spectral radius. Standard definitions of these metrics can be found in ref. (Geng Li, 2012).

Note that the feature extraction process is performed once for each cohort and cell type of interest, and it can be used multiple times for subsequent downstream analysis tasks.

#### Data visualization and exploration

Once the feature matrix has been calculated, it can be fed to the TIMEClust program. TIMEClust deploys an unsupervised clustering approach to group together single cells based on their neighbourhood’s phenotypic, functional, and structural similarity. The consensus clustering method from the Spectre package (Ashhurst, et al., 2022) is used to identify a statistically robust number of clusters, without manual tuning. Based on this unbiased stratification of cellular neighbourhoods, LOCATOR offers several data exploration and visualization utilities. Users can generate and export box plots and bar plots to visualize the abundance of clustered cells, and subsequently produce summary heatmaps of the feature matrices (using standard R packages such as ggplot and pheatmap). Using LOCATOR’s API users can also perform the t-distributed stochastic neighbour embedding (t-SNE), a state-of-the-art statistical method (Maaten, L. and Hinton, G., 2008) for visualizing multidimensional cellular clusters into the 2D space. The Rtsne implementation has been incorporated into our R software implementation. Single cell clusters can be also mapped to their original tissue coordinates providing as an output high-resolution tissue images for data inspection and exploration.

Furthermore, *TIMEClust* offers utilities for TIME-driven exploration at the patient/cohort level. For each cellular cluster, we compute the abundance of clustered cells per patient. Then, the average/median abundance per cluster at the cohort level is used as a cut-off to identify individual patients that are enriched or depleted for the corresponding cellular cluster. Supplementary Figure 7 shows examples of tissue images that are enriched and depleted for specific cellular clusters. At the patient level, this simple quantification can be used as a basis for data-driven patient stratification and association with clinical and biological annotations. The LOCATOR software provides univariate survival analysis based on Kaplan-Meier method (survminer R package), and survival plots are automatically generated for every cellular cluster. A list of Alluvial plots (alluvial R package) is also produced to visualize cellular clusters and their association with clinicopathological information. Command-line syntax for running *ExtractFeatures* and *TIMEClust* programs is provided on our GitHub page. More detailed information about the dependent R packages and examples/tutorials are also available.

### Breast cancer cohorts and clinicopathological information used in this study

For the development and evaluation of LOCATOR we use publicly available MCI data from Keren et al. (Keren, et al., 2018) and Danenberg et al. (Danenberg, et al., 2022) herein we call them as “Keren dataset” and “Danenberg dataset” respectively. In the first dataset/study, Keren et al. used multiplexed ion beam imaging by time-of-flight (MIBI-TOF) to quantify 36 markers in 41 triple negative breast cancer (TNBC) patients. The authors deployed an algorithm to investigate how immune and tumour cells interact with each other in respect to immune cell interactions. Using the deployed algorithm, TIME architectures were annotated as Mixed and Compartmentalized tissue types and the authors validated that the Compartmentalized type was associated with improved survival. The authors also explored the expression of PDL1 immunoregulatory protein, in association to the spatial structure of the Compartmentalized and Mixed tissue types.

In the second dataset/study, Danenberg at el. used Hyperion IMC to generate 37-dimensional images of 693 breast cancer tumours matched with genomic and clinical information. Using spatial tissue-based analyses, and clustering of spatial regions the authors identified ten TIME structures/clusters and explored how these clusters (IntClust information is available) were linked with patient-specific somatic alterations, ER status, tumour grade, and PAM50 breast cancer subgroups. Summary statistics about the datasets and patient-specific annotations used in both studies/datasets can be found in Supplementary Figure 2.

## Discussion

In this study we present the development and use of LOCATOR, a new computational tool to analyse highly multiplexed imaging data from cancer tissues. LOCATOR deploys a versatile graph-based representation of cellular neighbourhoods to engineer features reflecting cellular phenotype abundance, cellular connectivity, and cell to cell spatial relationships. Subsequently, it offers visualization utilities as well as a simple unsupervised clustering approach to stratify patients based on the spatial similarity of their cellular neighbourhoods. The tool is automated and modular, its programs are customizable by users, and they can be widely applied irrespective of the study design, the imaging technology used, and the available antibody panels. In this way, LOCATOR opens possibilities to standardize MCI downstream analyses contributing towards hypothesis testing and TIME-driven exploration of cellular microenvironments via MCI.

To test LOCATOR, we perform two case studies using publicly available data from two seminal studies in breast cancer. Since the knowledge about the role of macrophages for patient stratification is limited, we use LOCATOR to analyse cellular microenvironments with macrophages as target cell type. Focusing on the Keren dataset, our patient stratification results are in concordance with clinicopathological parameters and other computationally derived published results. This is also the case for a larger cohort of MCI samples (Danenberg dataset), where LOCATOR contributes an alternative way for patient stratification based on the spatial organisation of macrophages. When compared with two independent spatial analysis algorithms, DenVar and lisaClust, LOCATOR achieves superior performance using Keren dataset as benchmark. In addition, a self-consistency test with synthetic data demonstrates LOCATOR’s power to produce reliable and robust results when analysing unseen patient cohorts. LOCATOR runs fast (approximately 15 min for feature generation and downstream analyses for ∼500 samples) in a commodity computer (16GB RAM with Intel i5 CPU 1.7 GHz), without requiring significant programming knowledge. Thus, it opens interesting possibilities to complement existing patient-risk stratification methods and enhance data-driven biomarker discoveries in future studies.

Focusing on technical aspects, many technical improvements are feasible. For example, implementing a graphical-user interface (GUI) would make LOCATOR even more interactive and easier for users with no prior programming knowledge skills. We are working towards this direction, providing improved customization of LOCATOR’s utilities. In addition, incorporating state-of-the-art feature selection methods (Soufan, et al., 2015) to select the most important features of TIMEs, might be useful for more effective TIME-driven patient risk stratification tasks. In addition, other machine learning-based methods can be incorporated for biomarker discoveries (Tislevoll, et al., 2023). Furthermore, the proposed graph-based representation of cellular neighbourhoods can be subjected to more advanced graph algorithms and analytic platforms taking advantage of high-performance computing infrastructures.

In terms of data analyses, the results presented here focusing mainly on macrophages corroborate many clinically relevant characteristics of patients. However, not all the detected patient clusters reach statistically significant associations with clinicopathological parameters. This is also the case for many of our clustering analysis results focusing on T-cytotoxic cells from the Keren dataset. We argue that this is expected given the complexity of the underlying datasets, the size of the clusters and the performance of the unsupervised clustering approach we deployed. Thus, new methods are needed that can combine data from multiple studies, and LOCATOR provides a step in this direction. Here, both case studies presented highlight LOCATOR’s ability to test hypotheses and unbiasedly explore and stratify patients using features of their TIMEs’.

In summary, LOCATOR streamlines the application of tissue imaging technologies for TIME-driven patient stratification. The outcome of this work may be useful for cancer research studies focusing on cellular heterogeneity, and precision cancer medicine. LOCATOR’s graph-based representation can be easily extended to describe even more comprehensive molecular profiles from single-cell multimodal assays and spatial transcriptomics (e.g., 10X Visium). In conclusion, we believe that the use of LOCATOR could provide a more general bioinformatics framework to explore the spatial organization of cellular communities in the broad context of human diseases.

## Acknowledgements

The authors would like to thank Dr. Vladan Milosevic (UiB) who kindly read the revised version of the manuscript and provided valuable feedback.

## Author Contribution

R.E developed LOCATOR and analysed the data; L.A and I.J provided intellectual input and helped edit the manuscript; D.K and R.E wrote the manuscript; D.K. supervised the research.

