## Supplementary Material for "LOCATOR: feature extraction and spatial analysis of the cancer tissue microenvironment using mass cytometry imaging technologies"

**Supplementary Table 1.** Overview of relevant tools and algorithms for data-driven analyses of TIMEs**.**

| Name | Reference | Programming language | Brief description | Available as a standalone tool | Used for patient stratification |
|---|---|---|---|---|---|
| DenVar | Vu, et al., 2022 | Python | Threshold-free algorithm in which distance between pairs of patients is computed based on the probability density of a functional protein in their TIMEs. The distance matrix is used to classify patients into clusters, but the analysis is limited to the expression levels of a single protein. | + | + |
| LisaClust | Patrick, et al., 2023 | R | The algorithm utilizes local indicators of spatial associations (LISA function) and performs unsupervised clustering to characterize cell interactions between different cellular classes. | + | - |
| SPF | Seal, et al., 2022 | R | The algorithm utilizes the K-function and its variants, to explore inter-cell dependencies as a function of distances between cells. | + | - |
| Keren et al. | Keren, et al., 2018 | - | One of the first computational frameworks to decipher TIMEs organization focusing on cellular composition, spatial arrangement of cells, and regulatory-protein expression. A computational score was proposed to identify mixed and compartmentalized tissue types in breast cancer. | - | + |
| Danenberg et al. | Danenberg et al. 2022 | R | Computational analysis framework that utilized spatial clustering of cellular neighborhood. The authors identified ten TIME clusters in breast cancer, including quiescent vascularized stroma and other spatial cellular patterns associated with somatic | - | - |

| | | | | | |
|---|---|---|---|---|---|
| | | | alterations and genomic breast cancer subtypes. | | |
| Giotto | Dries, et al. 2021 | Python | A comprehensive and open-source toolbox for spatial data analysis and visualization. It implements a wide range of algorithms to characterize tissue composition, spatial expression patterns, and cellular interactions. Giotto can be applied to MCI but has a primarily focus on spatial transcriptomic data. | + | + |
| LOCATOR | Ehsani, et al. 2023 | R | An automated and open-source framework for the analysis of TIMEs. It proposes a graph-based representation of TIMEs to describe features of the cellular organisation and deploys downstream analysis and visualisation utilities that can be used for data-driven patient risk stratification. | + | + |

**Supplementary Table 2:** Summary results of the statistical tests (Fisher's exact test) we performed to associate cellular clustering findings with clinicopathological parameters.

| CluterType | ClinicalType | +/- | +/+ | -/- | -/+ | P value |
|---|---|---|---|---|---|---|
| NC2 | G3 | 0.24 | 0.76 | 0.46 | 0.54 | 0 |
| NC2 | ER+ | 0.37 | 0.63 | 0.15 | 0.85 | 0 |
| NC2 | ER- | 0.63 | 0.37 | 0.85 | 0.15 | 0 |
| NC2 | Basal | 0.76 | 0.24 | 0.88 | 0.12 | 0.001 |
| NC2 | LumA | 0.78 | 0.22 | 0.58 | 0.42 | 0 |
| NC2 | Her2 | 0.84 | 0.16 | 0.91 | 0.09 | 0.039 |
| NC2 | IntClust 7 | 0.95 | 0.05 | 0.87 | 0.13 | 0.007 |
| NC2 | IntClust 8 | 0.95 | 0.05 | 0.84 | 0.16 | 0 |
| NC2 | IntClust 10 | 0.83 | 0.17 | 0.93 | 0.07 | 0.001 |
| NC2 | IntClust 4- | 0.88 | 0.12 | 0.96 | 0.04 | 0.008 |
| NC3 | G3 | 0.5 | 0.5 | 0.32 | 0.68 | 0 |
| NC3 | G1 | 0.9 | 0.1 | 0.96 | 0.04 | 0.01 |
| NC3 | ER+ | 0.17 | 0.83 | 0.26 | 0.74 | 0.019 |
| NC3 | ER- | 0.83 | 0.17 | 0.74 | 0.26 | 0.019 |
| NC3 | NL | 0.87 | 0.13 | 0.93 | 0.07 | 0.042 |
| NC3 | LumB | 0.78 | 0.22 | 0.68 | 0.32 | 0.028 |
| NC8 | G3 | 0.07 | 0.93 | 0.42 | 0.58 | 0 |
| NC8 | ER+ | 0.43 | 0.57 | 0.21 | 0.79 | 0.002 |
| NC8 | ER- | 0.57 | 0.43 | 0.79 | 0.21 | 0.002 |
| NC8 | Basal | 0.97 | 0.03 | 0.83 | 0.17 | 0.019 |
| NC8 | LumA | 0.87 | 0.13 | 0.63 | 0.37 | 0.001 |
| NC8 | Her2 | 0.54 | 0.46 | 0.92 | 0.08 | 0 |
| NC8 | IntClust 7 | 1 | 0 | 0.88 | 0.12 | 0.024 |
| NC8 | IntClust 8 | 1 | 0 | 0.86 | 0.14 | 0.009 |
| NC8 | IntClust 5+ | 0.62 | 0.38 | 0.97 | 0.03 | 0 |
| NC8 | IntClust 3 | 1 | 0 | 0.86 | 0.14 | 0.009 |
| NC8 | IntClust 10 | 1 | 0 | 0.89 | 0.11 | 0.023 |
| NC8 | IntClust 5- | 0.59 | 0.41 | 0.97 | 0.03 | 0 |

**Supplementary Table 3:** Comparison analysis results between DenVar, LisaClust_modified and LOCATOR using as benchmark tissue annotations published by Keren et al. Performance metrics used are Accuracy (ACC), Sensitivity (SN) and Specificity (SP).

| **Input marker: PDL1** | | | | | | | | | | | | | |
|---|---|---|---|---|---|---|---|---|---|---|---|---|---|
| | **DenVar** | **LisaClust_modified** | | | | | | **LOCATOR** | | | | | |
| | | **NC1** | **NC2** | **NC3** | **NC4** | **NC5** | **Mean** | **NC1** | **NC2** | **NC3** | **NC4** | **NC5** | **Mean** |
| **ACC** | 0.62 | 0.57 | 0.67 | 0.80 | 0.83 | 0.83 | **0.74** | 0.87 | 0.77 | 0.52 | 0.65 | 0.81 | 0.72 |
| **SN** | **0.86** | 0.60 | 0.75 | 0.75 | 0.87 | 0.79 | 0.75 | 0.81 | 0.73 | 0.55 | 0.61 | 0.75 | 0.69 |
| **SP** | 0.56 | 0.53 | 0.61 | 0.90 | 0.80 | 0.91 | 0.75 | 1.0 | 0.89 | 0.46 | 1.0 | 0.87 | **0.84** |
| **Input marker: CD68** | | | | | | | | | | | | | |
| | **DenVar** | **LisaClust_modified** | | | | | | **LOCATOR** | | | | | |
| | | **NC1** | **NC2** | **NC3** | **NC4** | **NC5** | **Mean** | **NC1** | **NC2** | **NC3** | **NC4** | **NC5** | **Mean** |
| **ACC** | 0.59 | 0.66 | 0.72 | 0.53 | 0.56 | 0.69 | 0.63 | 0.88 | 0.55 | 0.91 | 0.64 | 0.58 | **0.71** |
| **SN** | 0.58 | 0.62 | 0.75 | 0.55 | 0.59 | 0.73 | 0.65 | 0.89 | 0.55 | 1.0 | 0.61 | 0.56 | **0.72** |
| **SP** | 0.67 | 0.75 | 0.69 | 0.50 | 0.53 | 0.65 | 0.62 | 0.87 | 0.50 | 0.83 | 0.80 | 1.0 | **0.80** |

**Supplementary Figure 1:** LOCATOR's workflow and pseudo code description of its' R programs.

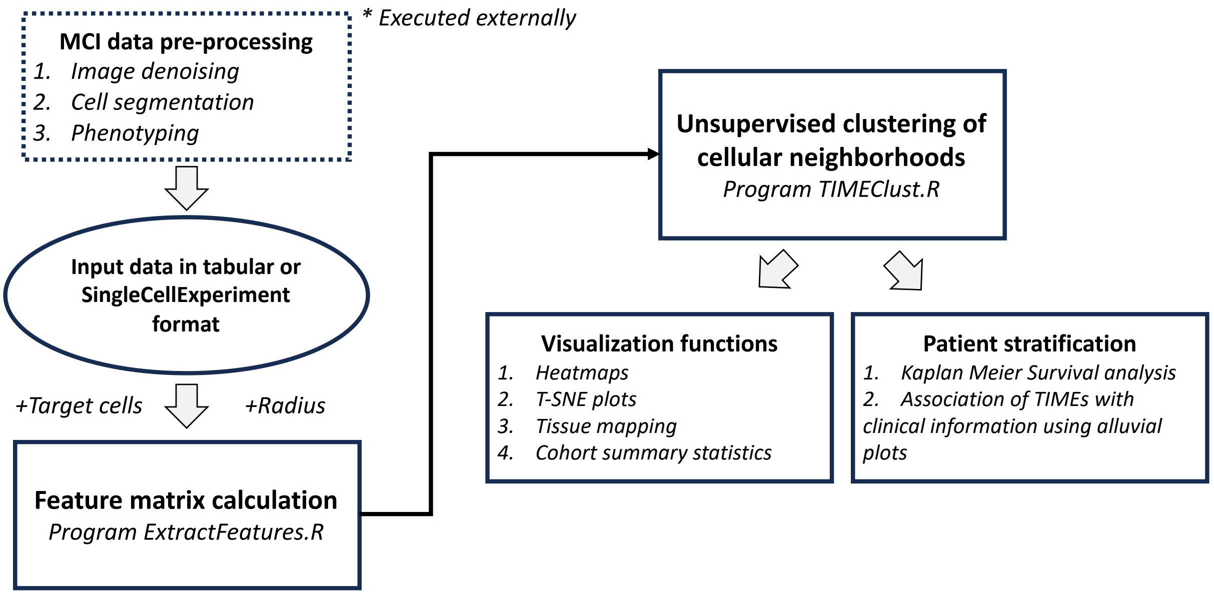


**Pseudocode for ExtractFeatures.R**

**Input:** pre-processed data in csv format or SingleCellExperiment, radius R, target cells (canonical cell type or cell subset)

**Columns/Variables used:** Target cell type, Meta cell type(tumor/non-tumor), Image ID, Cell ID, selected markers' expression, X and Y coordinates.

**Output:** Feature matrix M (numeric)

1: For each target cell {
2: Generate a circle C using target's cell $X_t$,$Y_t$ coordinates as center and R as radius.
3: Estimate cell type frequencies for all cells within C (CT features).
4: Generate an undirected graph G with cells as vertices and identify cell–cell interactions (edges) using Voronoi algorithm.
5: Estimate Connectivity score for all cell types from G (CS featurs)
6: Estimate graph theory metrics from G (CG features)
7: Concatenate feature vectors CT,CS, CG from step 3,5,6.
}
8: Return feature matrix M

**Pseudocode for TIMEClust.R**

**Input:** Feature matrix M from ExtractFeatures.R, clinical info (i.e., survival time, molecular type) per patient.

**Outputs:** a tabular file (i.e., csv) with the clustering results of the target cells; heatmaps, T-SNE plots, Tissue mapping, and summary statistics about the clustering results at the cohort level; survival analysis using the clustering results, alluvial plots combining clinical info and clustering results.

```
1: Unsupervised clustering of cellular neighborhoods (NCs) from M, by FlowSOM algorithm using
Spectre package.
2: Perform visualization of clustering results
        2a: Feature vector visualization as heatmap
        2b: 2D representation of clusters using T-SNE
        2c: Tissue mapping of clustering results per patient using X,Y cell coordinates and NC info
        2d: Summary statistics of clustering results at the cohort level
3: Perform survival and alluvial analysis of clustering results
        For each NC {
                3a: Estimate the abundance of NC per patient.
                3b: Identify the average/median abundance across all patients (cutoff)
                3c: Use cutoff from 3b to detect individual patients enriched or depleted for NC.
                3d: Perform univariate Kaplan-Meir survival analysis (depleted vs. enriched for NC).
                3e: Perform Alluvial analysis/plots (depleted vs. enriched for NC) using input clinical info
                and estimate statistical significance of proportions.
        }
4: Return visualization plots and clustering analysis results
```

**Supplementary Figure 2:** Summary statistics and annotations for Keren and Danenberg datasets.

a)

| Dataset | #sample | #markers | #cell | #cell types | #samples used in LOCATOR | Clinical Annotations |
|---|---|---|---|---|---|---|
| **Keren et al.** | 41(36*) | 36 | 190,953 | 11 | 33 | Mixing score<br>Survival |
| **Danenberg et al.** | 693 | 37 | 1,045,835 | 22 | 497 | PAM50<br>IntClust<br>Tumour Grade<br>ERStatus<br>Survival |

b)

| | Annotations | Distinguishing Features | Prognosis |
|---|---|---|---|
| **PAM50** | NL (Normal Like) | ER+, PR+, HER2- | Good |
| | LumA (Luminal A) | ER+, PR+\PR-, HER2- | Good |
| | LumB (Luminal B) | ER+, PR+\PR-, HER2+\ HER2- | Intermediate |
| | Her2 | ER-, PR-, HER2+ | Poor |
| | Basal | ER-, PR-, HER2- | Poor |
| **IntClust** | IntClust 1 | 17q23 amplification<br>GATA3 mutations<br>High genomic instability | Intermediate |
| | IntClust 2 | 11q13-14 amplification (CCND1)<br>High genomic instability | Poor |
| | IntClust 3 | Low genomic instability with few copy number changes<br>PIK3CA and CDH1 mutations | Good |
| | IntClust 4- | ER-negative tumors irrespective of HER2 status | Poor |
| | IntClust 4+ | Low genomic instability<br>Up regulation immune response genes | Good |
| | IntClust 5- | ERBB2 amplification | Poor |
| | IntClust 5+ | ERBB2 abbreviation | Good |
| | IntClust 6 | 8p12 amplification (ZNF703)<br>High genomic instability | Intermediate |
| | IntClus 7 | 16p gain, 16q loss, 8q amplification<br>MAP3K1 mutations | Good |
| | IntClust 8 | 1q gain, 16q loss<br>PIK3CA and GATA3 mutations | Good |
| | IntClust 9 | 8q gain, 20q amplification<br>High genomic instability<br>TP53 mutations | Intermediate |
| | IntClust 10 | 5q loss, 8q gain, 10p gain, 12p gain<br>Impaired DNA checkpoint regulation, TP53 mutations | Poor |
| **Grade** | G1 (Grade 1) | slower-growing and less likely to spread | Good |
| | G2 (Grade 2) | growing faster than a grade 1 cancer but slower than a grade 3 cancer | Intermediate |
| | G3 (Grade 3) | faster-growing cancer that's more likely to spread | Poor |
| **ER status** | ER + | Cancer cells have receptors that allow them to use the hormone estrogen to grow | Good |

c)

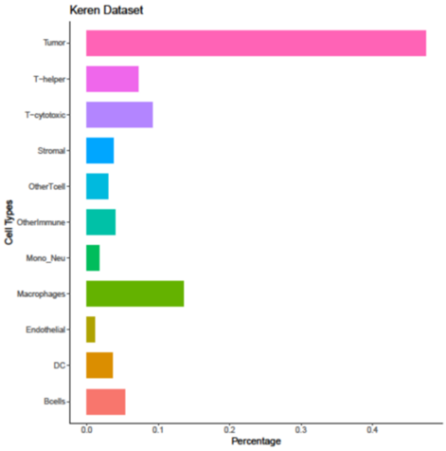


d)

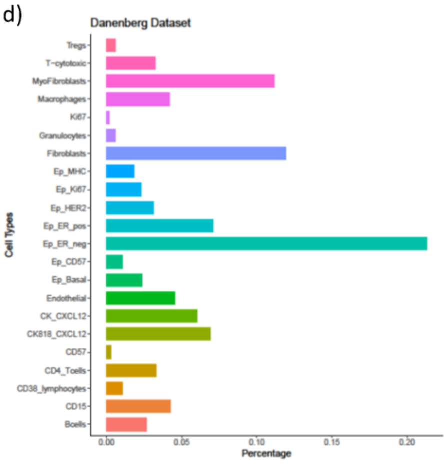

**Supplementary Figure 3: LOCATOR outputs for Keren dataset focusing on T cytotoxic cells.** a) Heatmap representation of the feature vector for the identified clusters. b) Bar plot showing cellular abundance of the identified cellular clusters. c) Boxplot showing the number of T-cytotoxic cells per patients across different cellular clusters. d)t-SNE visualisation of the identified cellular clusters. e) Example of tissue mapping with cellular cluster annotation from one patient (patient No 7). Gray colour indicates tumour cells and pink colour indicates non-tumour cells. All other colours in the image correspond to the detected cellular clusters from subplots c and d.

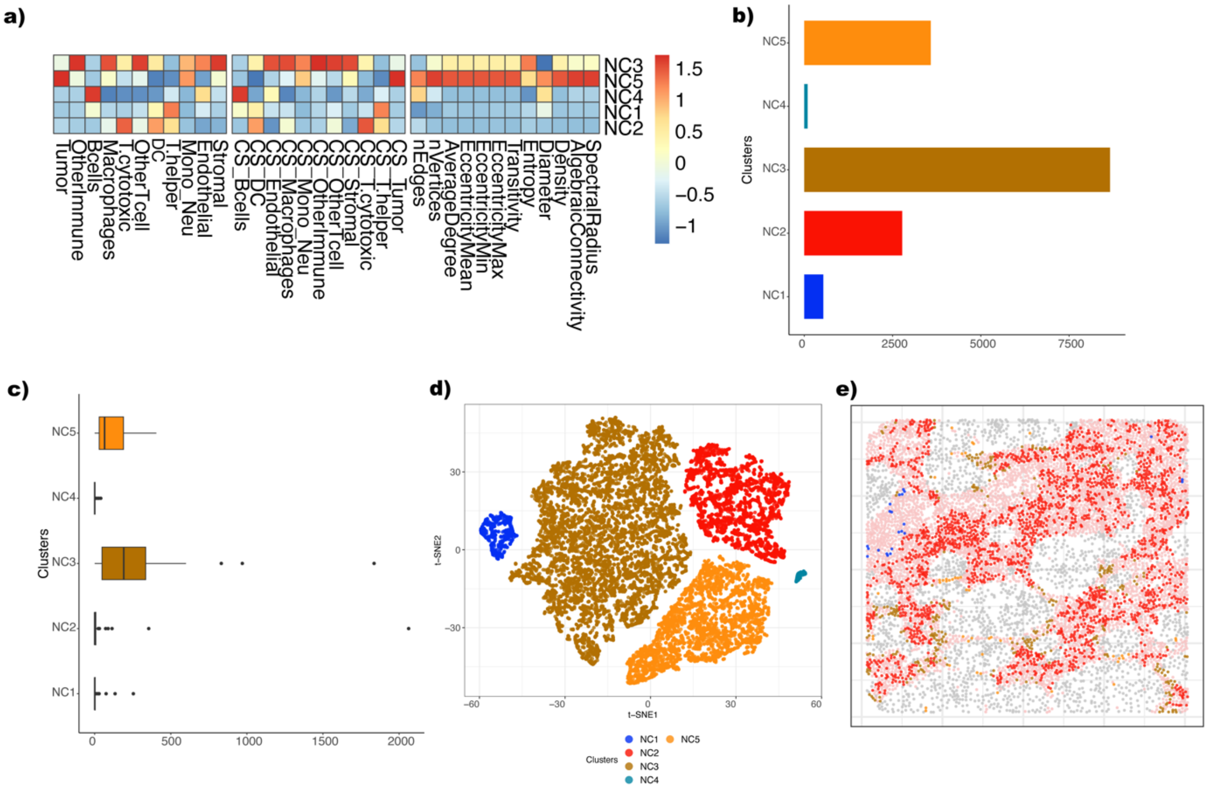

**Supplementary Figure 4: LOCATOR outputs for Danenberg dataset focusing on T cytotoxic cells.** a) Heatmap representation of the feature vector for the identified clusters. b) Bar plot showing cellular abundance of the identified cellular clusters. c) Boxplot showing the number of T-cytotoxic cells per patients across different cellular clusters. d)t-SNE visualisation of the identified cellular clusters. e) Example of tissue mapping with cellular cluster annotation from one patient (patient No 47). Gray colour indicates tumour cells and pink colour indicates non-tumour cells. All other colours in the image correspond to the detected cellular clusters from subplots c and d.

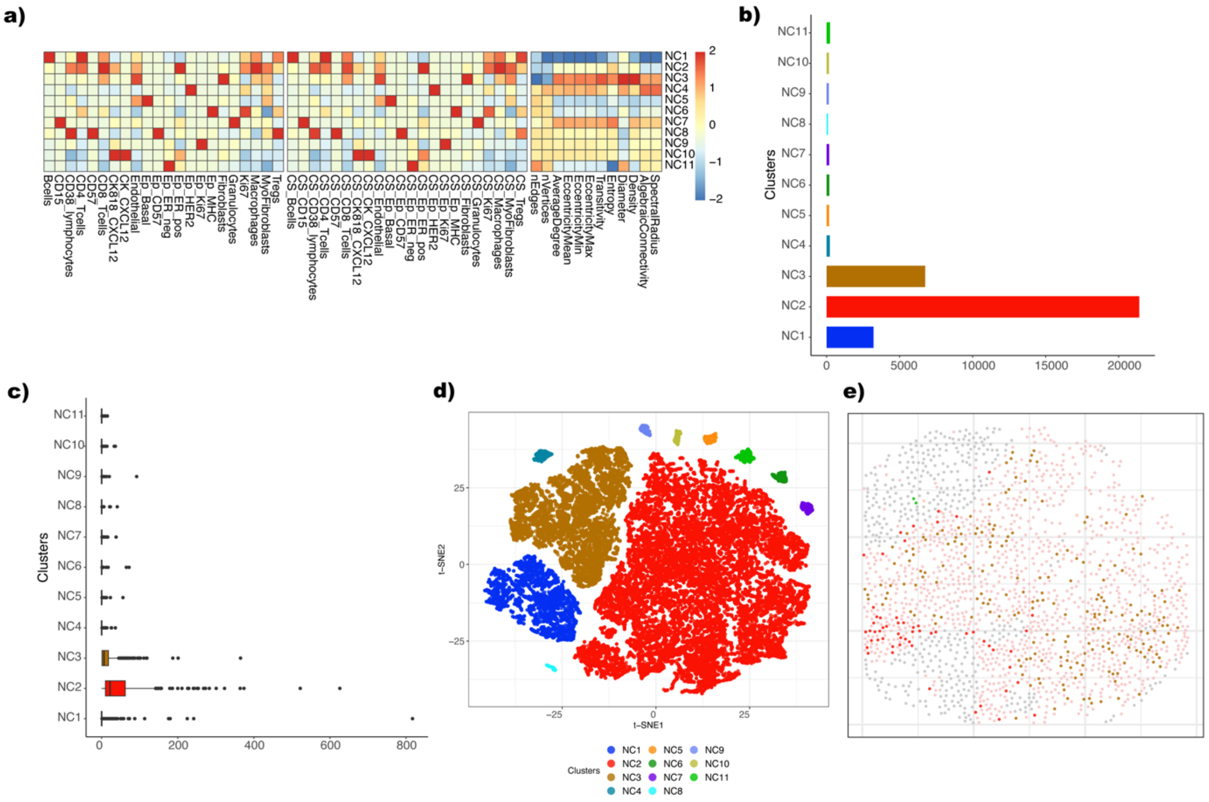

**Supplementary Figure 5: Case study 2: Association of NC2, NC3 and NC8 identified cellular cluster with clinicopathological parameters and independent published results.** a-i) Alluvial plots showing the fraction of NC2, NC3 and NC8 enriched/depleted patients and their underlying associations with histological Grade, IntClust results and ER status. Significance of the results is obtained using Fishers-exact test at a p-value level of 0.05 (results marked with *, and Supplementary Table 2).

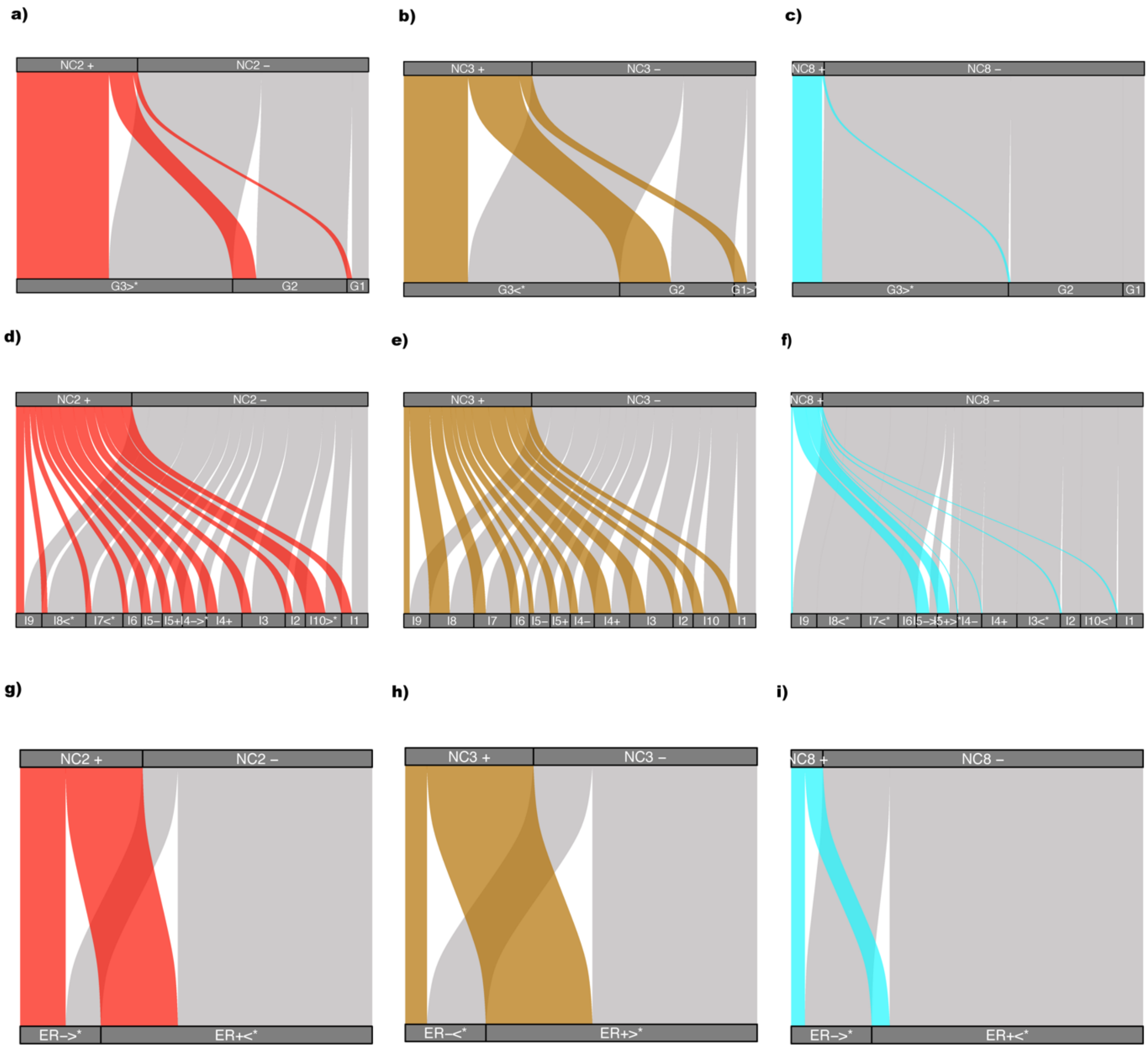

**Supplementary Figure 6: LOCATOR application on T-cytotoxic cells from Case study 2: Association of NC2, NC3 and NC4 identified cellular clusters with survival and other clinicopathological parameters.** a-c) Survival plots where patients are classified as enriched (NC2+, NC3+, NC4+) or depleted (NC2- , NC3-, NC4-) based on their mean cellular abundances. d-m) Alluvial plots showing the fraction of NC2, NC3 and NC4 enriched/depleted patients and their underlying PAM50 classification, histological grade and ER status.  Significance of the results is obtained using Fishers-exact test at a p-value levels of 0.05.

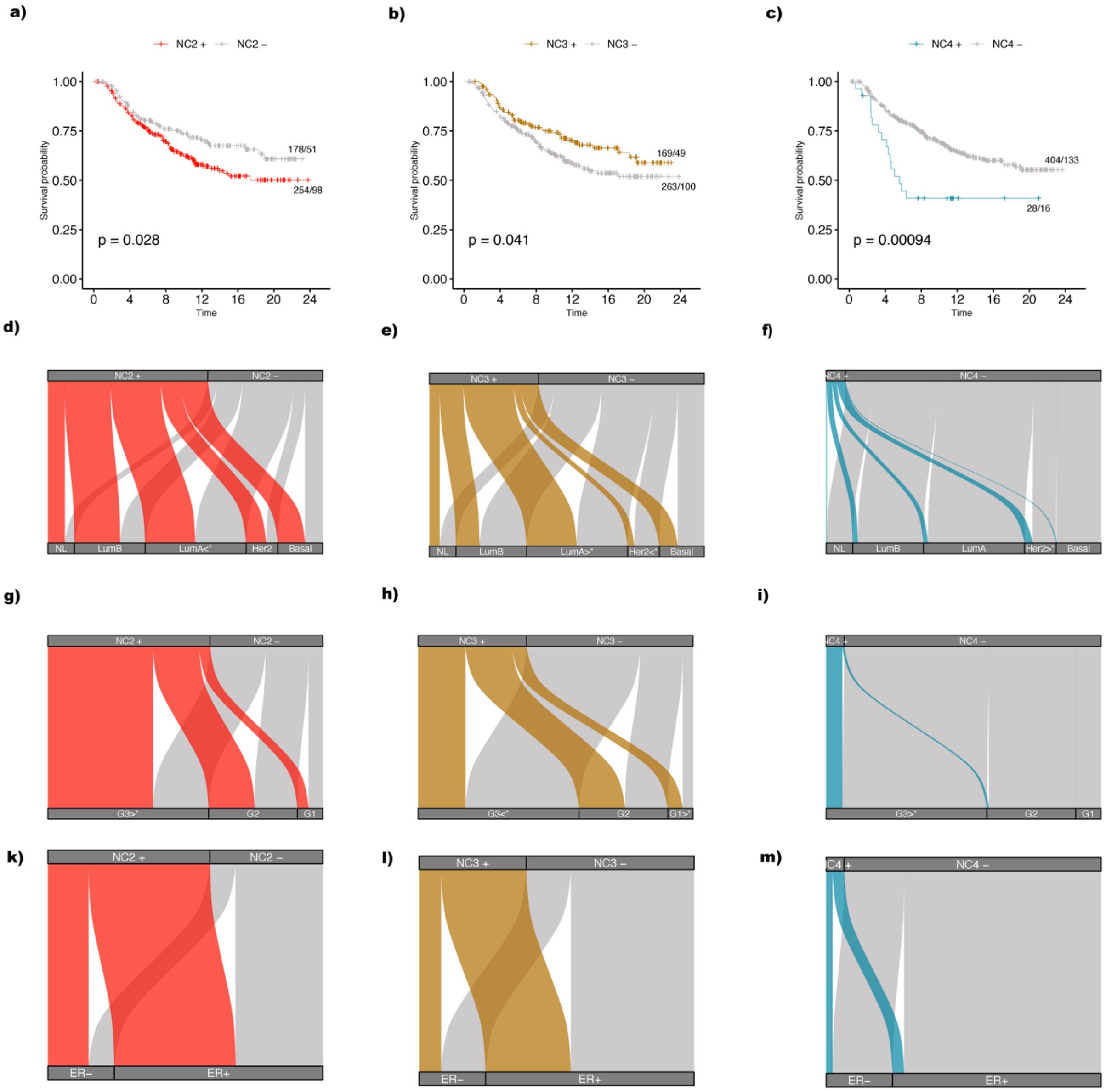

**Supplementary Figure 7: Examples of tissue reconstruction and mapping generated by LOCATOR visualisation utilities.** a-b) Selected tissue images from Keren dataset. c-d) Selected tissue images from Danenberg dataset. In all subplots, gray colour indicates tumour cells and pink colour indicates non-tumour cells. All other colours correspond to cellular cluster annotations reported in Figure 2 and Figure 4.

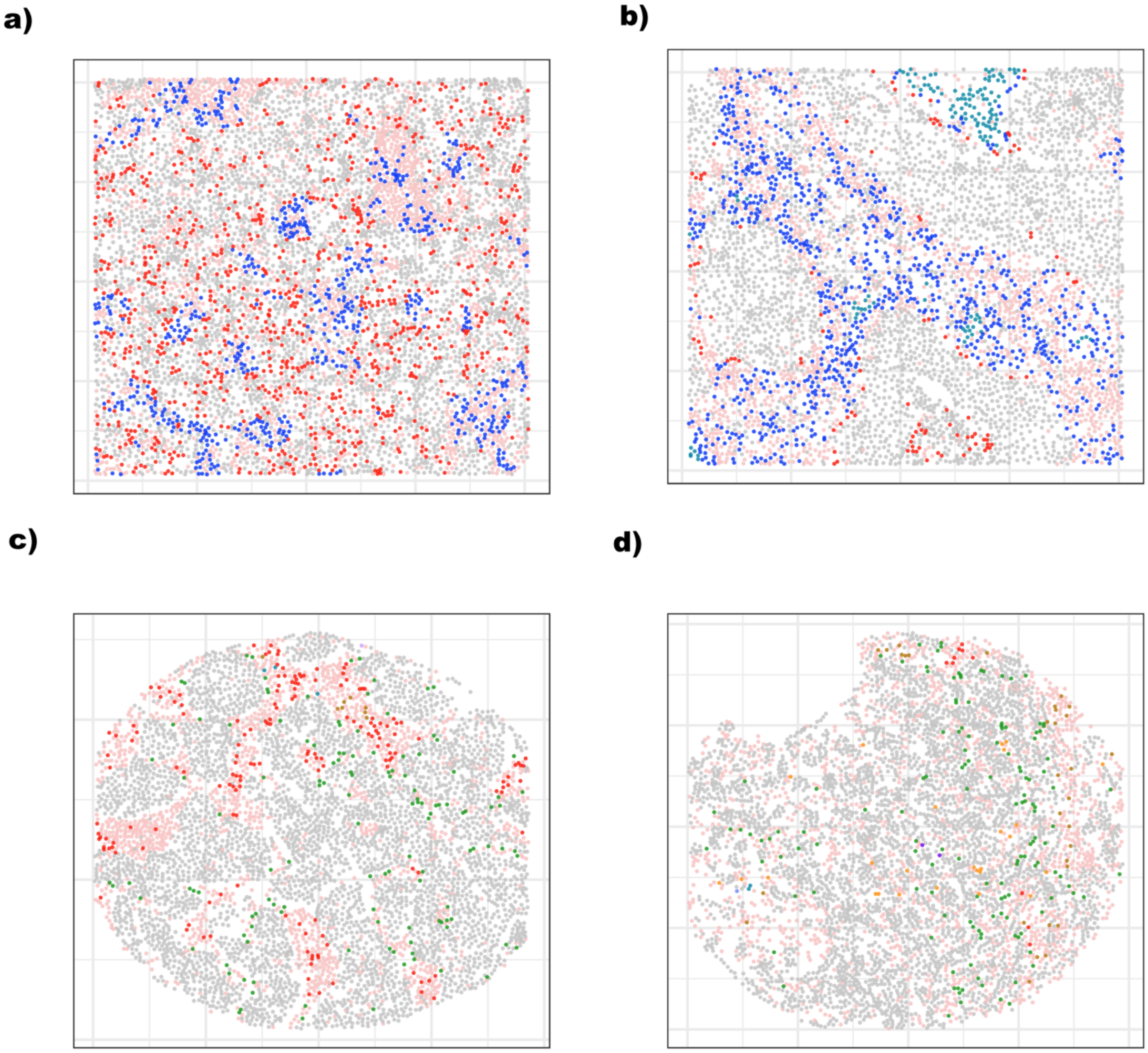
